# Active sensing movements by antennae enhance neural processing and plasticity for odour processing in the CNS in the honey bee *Apis mellifera*

**DOI:** 10.64898/2026.09.14.751598

**Authors:** Nisha Kulkarni, Harry H Lei, Jake Leyko, Brian H Smith, Hong Lei

## Abstract

Animals frequently move their sensory organs relative to dynamic environmental stimuli to maximize information acquisition, a behaviour known as active sensing. Recent studies show that insects adjust antennal movements according to the temporal structure and salience of olfactory signals, suggesting that antennae participate in active sensing. However, how these movements affect central olfactory processing remains unclear. To address this question, we trained honey bee foragers using a proboscis extension response (PER) protocol in which one odour (CS+) was paired with sucrose and another (CS−) with salt water. This training produces distinct antennal movements toward the two odours. Bees were subsequently tested with a series of concentrations of both CS+ and CS− odours. Antennae were either fixed or allowed to move freely during training and testing, while electrophysiological recordings were made from the antennae and from the antennal lobes. Peripheral antennal responses across odour concentrations were not affected by whether the antennae were fixed or free. In contrast, antennal lobe projection neurons and local interneurons showed learning-related increases in responses to the CS+ odour, consistent with previous studies, and these increases were stronger in bees with freely moving antennae. Neural responses were also more variable, particularly at lower odour concentrations, when antennae could move, suggesting that response variability may contribute to plasticity. These results support the hypothesis that active antennal movements influence central processing of odour representations. Future studies comparing restrained bees with freely walking bees will help clarify the natural behavioural contexts in which antennal movements enhance olfactory sensing.

**SUMMARY STATEMENT:** This study shows that allowing honey bees to move their antennae enhances learning-related neural plasticity, linking active sensing behaviour to central olfactory processing.

## INTRODUCTION

All animals rely on sensory systems to detect and encode environmental stimuli. They employ different behavioural strategies to expose sensors to various sources of information. Active sensing involves sensor movement to efficiently acquire information from the environment. Sensor movement can be achieved by adjusting body movement or by movement of the sensory structures (Zweifel and Hartmann, 2020). Classical examples of active sensing include echolocation in bats or electrolocation in fishes because they actively produce sound or electric waves, then gather information from the returning echoes. In olfaction in mammals, sniffing frequency (Spencer et al., 2021; Thesen et al., 1993; Wachowiak, 2011), nose motion (Khan et al., 2012), and body movements (Biswas et al., 2018) can be changed to improve sensing. As an analogous example in insects, active sensing includes antenna movements (Cholé et al., 2022; Claverie et al., 2023; Hoffmann and Couzin-Fuchs, 2023; Lei et al., 2022; Suver et al., 2023), the casting-and-surging behaviour in flying insects (Cardé, 2021; Murlis et al., 1992), and insect wing motion that allows improved sampling (Sane and Jacobson, 2006).

Several lines of argument imply that insect antennal movements likely represent active sensing of odours. First, antennal movement differs upon odour stimulation versus air stimulation. In honey bees, antennae move laterally, across the air flow, when a novel odour is presented (Birgiolas et al., 2017; Lei et al., 2022). Locusts increase antennal sweep frequency, reduce sweep amplitude, and move antenna towards an odour source (Huston et al., 2015). Cockroaches shift antennae toward an odour source, increase vertical movement and begin high-frequency local oscillations (Hoffmann and Couzin-Fuchs, 2023). Similarly, in bumble bees a low-frequency, large-amplitude oscillation of the antennae is triggered by the presence of odour stimuli. Moreover, the presence of an odour induce the antennae to oscillate vertically with a larger amplitude and laterally with a smaller amplitude, forming a cone of sampled airspace (Claverie et al., 2023). Second, antennal movement is modified based on stimulus valence. Cholé and colleagues tested how honey bee antennal movement is affected by odours with different innate biological values (Cholé et al., 2022). They found that antennae moved faster and moved toward sources of octanoic acid (royal jelly), methyl linoleate (brood pheromone), the mandibular queen pheromone, and the floral compound octanal. The antennae moved away from and slower to 2-heptanone (alarm pheromone). Similarly, Lei et al. reported that honey bee antennae moved towards novel odours and moved away from repeated non-rewarded familiar odours (Lei et al., 2022). In a recent study in cockroaches, a weaker antennal responses and slight increase in walking speed were found in response to a low valence odour (linalool) (Hoffmann and Couzin-Fuchs, 2023). Third, antennal movements exert impact on sensory processing either at detection level or coding level or both. In fruit flies, Suver and colleagues optogenetically activated specific antennal muscle motor neurons, which positioned the third segment of the antenna in such a way that the difference of antennal displacement due to wind produces a higher gain and acuity in encoding wind direction (Suver et al., 2023). Huston et al. reported that when a locust encounters an odour, it increases its antennal sweep frequency and shifts its sweep elevation to a point closer to the odour.

In spite of the evidence that antennal movements represent active sensing, it is not well understood how these movements affect neural activity in the CNS. Here we evaluate whether and how movements influence neural processing in the first synaptic relay for olfactory sensory information in the antennal lobes of the honey bee (*Apis mellifera*) brain. We differentially conditioned two treatment groups of bees to odours using a Proboscis Extension Response differential conditioning protocol (Smith and Burden, 2014), where one odour was associated with sucrose reinforcement (CS+) and the other with salt punishment (CS-). One group had antennae fixed in place, preventing movement, and the other group had antennae left free to move. The effects of antennal treatment (fixed vs. free) were assessed physiologically with neural recordings from the antennae and from the antennal lobes. We show that free antennal movement results in increased firing rate in local interneurons and projection neurons, and that these responses engage mechanisms of plasticity previously shown to be important for odour processing in the antennal lobes (Jernigan et al., 2020; Latshaw et al., 2023). These results suggest that active sensing movements of honey bee antennae are essential for fine tuning neural processing and, potentially, decision making in the brain.

## MATERIALS AND METHODS

### Animal preparation

Honey bees were prepared following a similar procedure described previously (Smith and Burden, 2014). In short, non-pollen forager honey bees (*Apis mellifera L*.) were collected in the morning at the entrance of a hive maintained outdoors in natural lighting conditions and brought into the lab. Bees were then briefly cooled restrained on a plastic stage with dental wax surrounding the head to keep it in place. After recovering from cooling, bees were then fed 2 μl of a 1.0 M sucrose solution until satiation and allowed to adjust to stages for approximately 1 hour in ambient conditions. Before recording, the antennae were immobilized with eicosane gently applied to the base of the antennae and the joint between flagellum and pedicel. A rectangular window was cut open on the cuticle between the compound eyes to expose the antennal lobes.

### Behavioural protocol

Honey bees were subjected to a differential learning protocol that combined the proboscis extension reflex (PER) with the presentation of two initially-neutral odours, 1-hexanol and 2-octanone. As described by Hadar and Menzel (Hadar and Menzel, 2010), the differential learning process conditions honeybees to associate neutral odours with either positive or negative stimuli. The PER was used as an indicator of successful associative learning, where the extension of the proboscis signified that the bee had learned to associate the odour with the reward.

The experimental procedure consisted of three phases: pre-training, training, and post-training. During the pre-training and post-training phases, a series of odour dilutions were presented to the honey bees for 0.5 seconds at 60-second intervals, with the dilutions ranging from 1:1000 to 1:1 for the fixed-antenna experiments and 1:100 to 10:1 for the free-antenna setup. The bees’ PER responses were observed to gauge their baseline behaviour toward the odours.

In the training phase, only the 10^-1^ concentration of the odours was used. One odour was assigned as positive, and the other as negative, with each presentation lasting four seconds. The two odours were presented in a pseudo-randomized order (ABBABAABAB with A and B being positive and negative odour, respectively). On the third second of each 4-second presentation, bees were either rewarded with sugar water for the positive odour or punished with salt water for the negative odour. This training phase was repeated over ten trials, with each bee receiving alternating positive and negative odour stimuli to reinforce the learning process.

For the free antennae recordings, the eicosane was removed from both the base of the antennae and the joint between the flagellum and pedicel prior to the differential learning protocol to allow for a full range of motion for the antennae. For the fixed antennae recordings, the differential learning procedure was administered without removing the eicosane from the antennae.

### Electroantennogram recordings

Electroantennogram (EAG) experiments followed a similar preparation process including the collection and staging followed by immobilization of the antennae. A thin silver reference electrode was inserted into the bee’s compound eye to provide a stable reference signal. A thin-walled glass capillary tube, filled with 0.5M LiCl solution, was used as the recording electrode. The antenna was gently inserted into the tip of the capillary tube, which had been coated with ultrasound transmission gel to ensure consistent contact between the antenna and the recording solution. This setup allowed for stable recordings of the electrical activity from the antenna.

Once the EAG setup was completed, the differential training protocol commenced, with odour stimuli being presented to the bees at specific intervals, as described in the behavioural protocol section. The EAG responses were recorded and analyzed to assess the bees’ sensory responses to different odour concentrations and associations

### Electrolobogram recordings

Electrolobogram (ELG) experiments followed a similar preparation process as the EAG method but with a few key modifications. First, eicosane was only applied to the bases of the antennae to largely limit their backward sweeping motions. Second, the bee’s head cavity was carefully dissected under a stereoscope to expose the brain. Trachea and connective tissues were removed with forceps to allow access to the antennal lobes. A silver reference wire was then inserted into the bee’s compound eye. Third, once a recording electrode was secured in the antennal lobe, the eicosane at the base was first kept to achieve fixed condition and then gently removed to achieve free condition. In this way, ELG signal was obtained from the same bee under both conditions.

Glass electrodes were used to record ELG. A thin-walled glass capillary tube was prepared using a P-2000 laser puller (Sutter® Instrument, CA) loaded with the manufacture-recommended whole cell recording parameters. The electrodes were then filled with saline by capillary action. The electrode was gently inserted into the brain region where antennal nerve enters the antennal lobe, and once a proper signal was observed, Kwiksil® (World Precision Instruments®, FL), a translucent medium-viscosity silicone adhesive, was applied to the cavity to secure the capillary tube in place and prevent brain dehydration, ensuring a stable connection to the antennal lobe.

### Multichannel extracellular recordings

Extracellular recordings were performed in the left antennal lobe with a 16-channel probe (NeuroNexus®, Ann Arbor, MI). Spike waveforms were digitized with a RZ2 system at a sampling rate of 20 KHz (Tucker-Davis Technologies®, Alachua, FL). In total, we recorded 115 units from 7 bees with fixed antennae and 75 units from 8 bees with free antennae. The fixed and free-antenna conditions were achieved using the same method as in the ELG experiment except that these two conditions were not manipulated on the same bee, considering the training protocol. The recording procedure followed previous publications with slight modifications (Lei et al., 2022). In short, after achieving stable recordings, Kwiksil® was applied into the brain cavity. Kwiksil® was used to prevent brain dehydration and also ensure stability of the probe. Honey bees were subjected to a differential learning procedure involving the proboscis extension reflex (PER). The bees were exposed to a series of odour/mineral oil dilutions (1:1000, 1:100, 1:10 and 1:1 v/v) with each dilution puffed for 0.5 seconds at 60-second intervals. The bee’s PER responses were manually noted.

During the training phase, two odours, 1-hexanol and 2-octanone, were used as neutral stimuli, with one odour assigned as positive (paired with a sugar water reward) and the other as negative (paired with a salt water punishment). The odours were presented to the bees for 4 seconds each, with the reward or punishment starting at the third second. This training process was repeated over ten trials, and the order of odour presentations was randomized. The differential learning procedure allowed the bees to associate each odour with either a positive or negative stimulus, as demonstrated by the PER.

### EAG/ELG data acquisition and analysis

The analog signals from the electroantennogram (EAG) and electrolobogram (ELG) recordings were amplified using an AxonClamp® 2B amplifier (Molecular Devices®, CA). These signals were digitized at a sampling rate of 2 kHz with a NI USB 6341 DAQ device and recorded using the NI DAQExpress© software (National Instruments®, TX). The recorded data were then exported into .csv format for further processing. For analysis, the data were imported into MATLAB© (MathWorks®, Natick, MA), where custom-written scripts were used to measure response magnitude.

### Analysis of extracellular recording data

From Synapse – Neurophysiology Suite of the TDT system, raw spike waveforms were exported to Offline Sorter© (Plexon® Inc, Dallas, TX) for classification into distinct clusters, representing individual units. This sorting was performed by projecting the waveforms onto a 3D space defined by the first three principal components, which captured the highest variance in waveform shapes. The tetrode configuration of the probe—where each block of recording sites contained four electrodes—allowed for enhanced discrimination of individual units. After classification, clusters were visually inspected and adjusted as necessary within the Offline Sorter software.

Once spike sorting was complete, timestamps of the sorted spikes were exported to NeuroExplorer© (Nex Technologies, Colorado Springs, CO) and MATLAB for further analysis. We used time-based instantaneous firing rate as the response metric, i.e. number of spikes divided by the time duration between the first and last spike within a predefined response window. The response window was dynamically determined in each experiment, considering the onset and offset of odour presentation, plus the response delay. Whereas the onset and offset of odour pulses were time-stamped by the acquisition system, the response delay was visually determined, thus different in each experiment. In short, we first generated a raster plot, peristimulus time histograms (psth) and a heatmap matrix across all units and all odour pulses in each experiment (Supplemental figure 1), then used the findchangepts function in MATLAB that automatically identifies points in the smoothed psth curve where statistical properties such as mean, variance, or trend change significantly to determine the start and end of the response, aided with visual correction if needed. This response window was then used universally in all units and all odours in this experiment.

Units were classified into putative projection neurons (PNs) and putative local neurons (LNs) using the statistical methods outlined in previous publications (Lei et al., 2022; Meyer et al., 2013). In short, we use four features in the classification: 1. Change in firing rate; 2. Lag to response; 3. Variability in the inter-spike interval; 4. Variability across odour trials. These four features were then used in a hierarchical clustering analysis using the Ward linkage method and the Euclidean distance metric (calculations were carried out with scipy’s linkage and fcluster functions). In total, we obtained 68 PNs and 44 LNs in the fixed-antennae group, 44 PNs and 31 LNs in the free-antennae group.

## RESULTS

### Effectiveness of training protocol and effects on peripheral responses

We performed differential training on two groups of bees, a fixed-antennae group (n=39) and a free-antennae group (n=20) (Supplemental figure 2A). Each group went through 5 training trials with CS+ (i.e. sugar rewarded odour) and 5 trials with CS- (i.e. salt solution punishment) in a pseudo-randomized order (Methods). In the fixed-antennae group, the proportion of bees showing responses, i.e. proboscis extension reflex (PER), increased significantly over the training trials to the CS+ odour (Kruskal-Wallis test, df=4, Chi-sq=52.73, p<0.0001), from 18% on the 1^st^ trial to 82% on the 5^th^ trial. In contrast, the proportion of bees responding to the CS-was about 20% on the 1^st^ trial and 0% on the 5^th^ trial, which was not significant (Kruskal-Wallis test, d.f.=4, Chi-sq=0.12, p=0.0871). The contrasting behavioural responses to the CS+ and CS-was also observed in the free-antennae group, demonstrating the effectiveness of the training protocol in both groups of bees. However, there was no significant difference between these two groups in differentiating between CS+ and CS-. For example, the percentage of differences between CS+ and CS-for both fixed and free antenna groups were nearly identical at 60% on trial 4 and about 80% on trial 5 (Supplemental figure 2B). Thus antennal restriction did not block or otherwise affect behavioural responses in this protocol, but we still help open the possibility that the restriction could affect upstream neural responses (see Discussion).

The first processing stage that could be affected by active sensing movements is with sensory transduction in olfactory sensory neurons of the antennae. Normally, one would measure field potentials via electroantennogram (EAG) recordings to evaluate antennal responses. However, due to the technical difficulties of recording EAGs from freely moving antennae – e.g. movement artifacts, displacement of electrodes - we recorded field potentials directly from the antennal lobe in the brain (electrolobogram; ELG) in the same individuals when antennae were first fixed and then let free. We first confirmed that the ELG signal is very similar to a conventional EAG signal, featuring a depolarization phase shortly after the onset of odour stimulation, then followed by a repolarization phase with much slower time constant (Methods and Fig. 1AB). Thus, the ELG recordings reflect sensory input to the brain from the antennae.

**Fig. 1.**
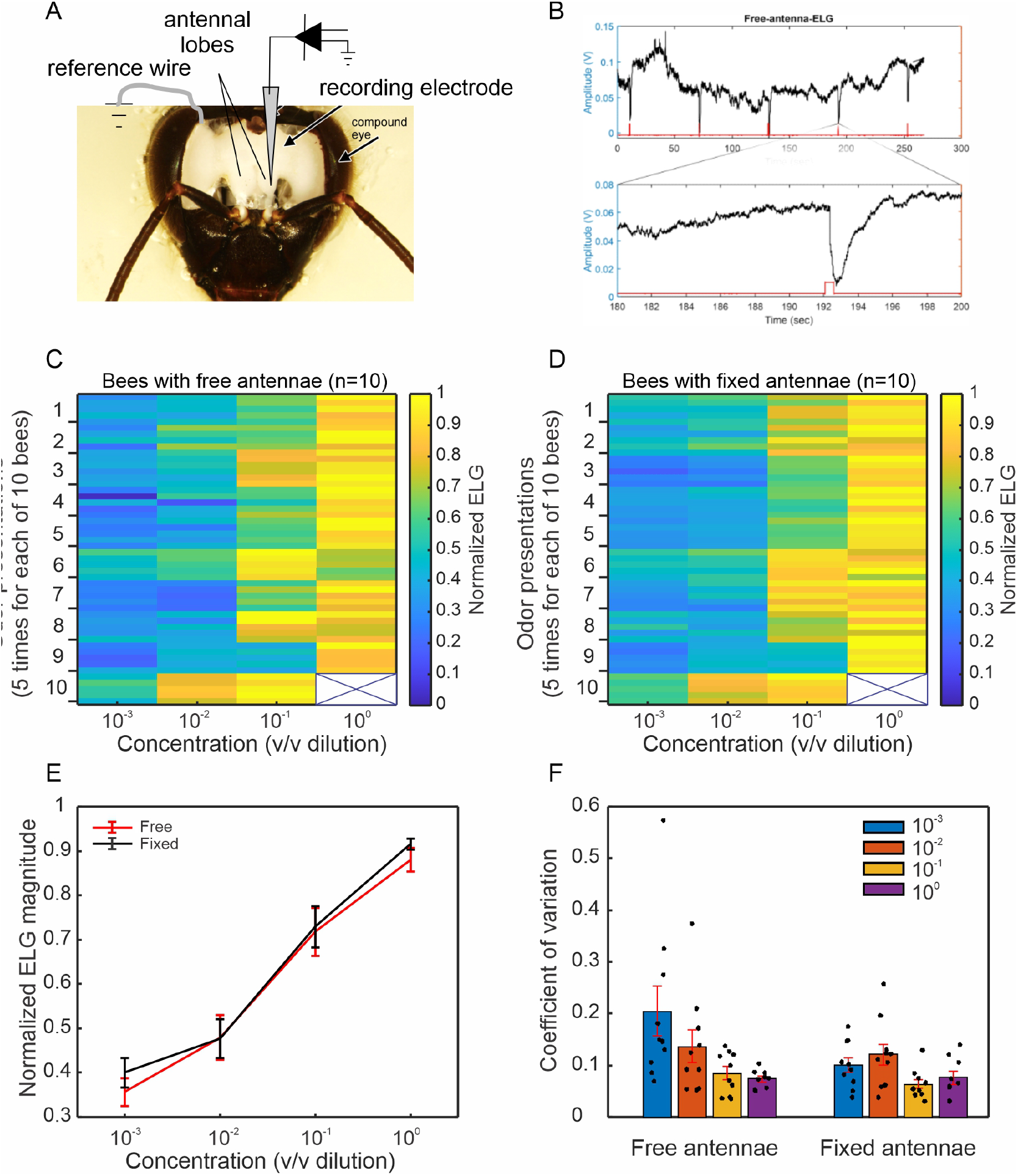
Free antennae promote response variation. A. Electrolobegram (ELG, see Methods) setup where the recording electrode was inserted into antennal lobe and the reference wire in compound eye. B. ELG trances showing odour-evoked responses to five odour presentations. The zoom-in view of a response (lower panel) shows that the ELG signal is highly similar to conventional electroantennogram (EAG) signals. C-D. The response magnitude in each bee is normalized to the maximal response in all odour presentations (2-octanone) and all concentrations (10^-3^, 10^-2^, 10^-1^ and 10^0^ vol/vol dilutions) in this bee, and then responses from all bees and all odour presentations are represented by the pseudo-colored matrices (C, bees with free antennae (n=10); D, bees with fixed antennae (n=10)). Bee #10 had missing data at the 10^0^ concentration. E. Bees with free (red line) and fixed (black line) antennae increased responses (mean ± s.e.) almost linearly following the increased odour concentrations, again validating the ELG signals. F. Coefficients of variation (CV, mean ± s.e.) to consecutive odour presentations at each concentration in free-antennae bees were generally higher than that in the fixed-antennae bees. Moreover, the CV decreased in the free-antennae group in a concentration-dependent manner. Such concentration dependency was lacking in the fixed antennae group.

When tested across a range of concentrations presented in a randomized sequence, the ELG data showed a strong concentration-dependent response in both fixed and free antennae treatments (Fig. 1C,D,E), which is consistent with EAG analyses. The means of these responses did not differ by antennal treatment (2-way ANOVA, concentration effect: p=1.56e-5; antennal treatment effect: p=0.26; interaction: p=0.99). However, free antennae produced more variable responses to consecutive odour stimulations in a concentration-dependent manner. Each animal was tested 5 times with each odour concentration. Inspection of the responses in the heat maps suggests that there is more high and low color variation across the trials within animals, particularly at the lower concentrations, when antennae were free (Fig. 1C) vs fixed (Fig. 1D). Furthermore, the coefficients of variation at the lower test concentrations were generally higher in the free-antennae group than in the fixed-antennae group (Fig.1F).

### Effects of active sensing on central plasticity

Next, we evaluated how antennal movement might facilitate odour processing downstream of the antennae, i.e. the primary olfactory center in insect antennal lobes. To do that we performed extracellular recordings from the antennal lobe while they were subjected to the same differential training protocol as described earlier. The bee’s antennae were either fixed or free. Both fixed- (n=7) and free-antenna (n=8) groups learned to discriminate the CS+ and CS-odours equally well, as previously. We recorded 115 units from 7 bees with fixed antennae and 75 units from 8 bees with free antennae. Based on their spontaneous firing patterns and response types (see Methods), these units were classified into putative PNs (n=68 in the fixed group; n=44 in the free group) and LNs (n=44 in the fixed group; n=31 in the free group) (Lei et al., 2022; Meyer et al., 2013). Generally, PNs exhibited significantly higher firing rates than LNs (Supplemental fig.3; Fig.2), further validating the statistical power of their separation.

**Fig. 2.**
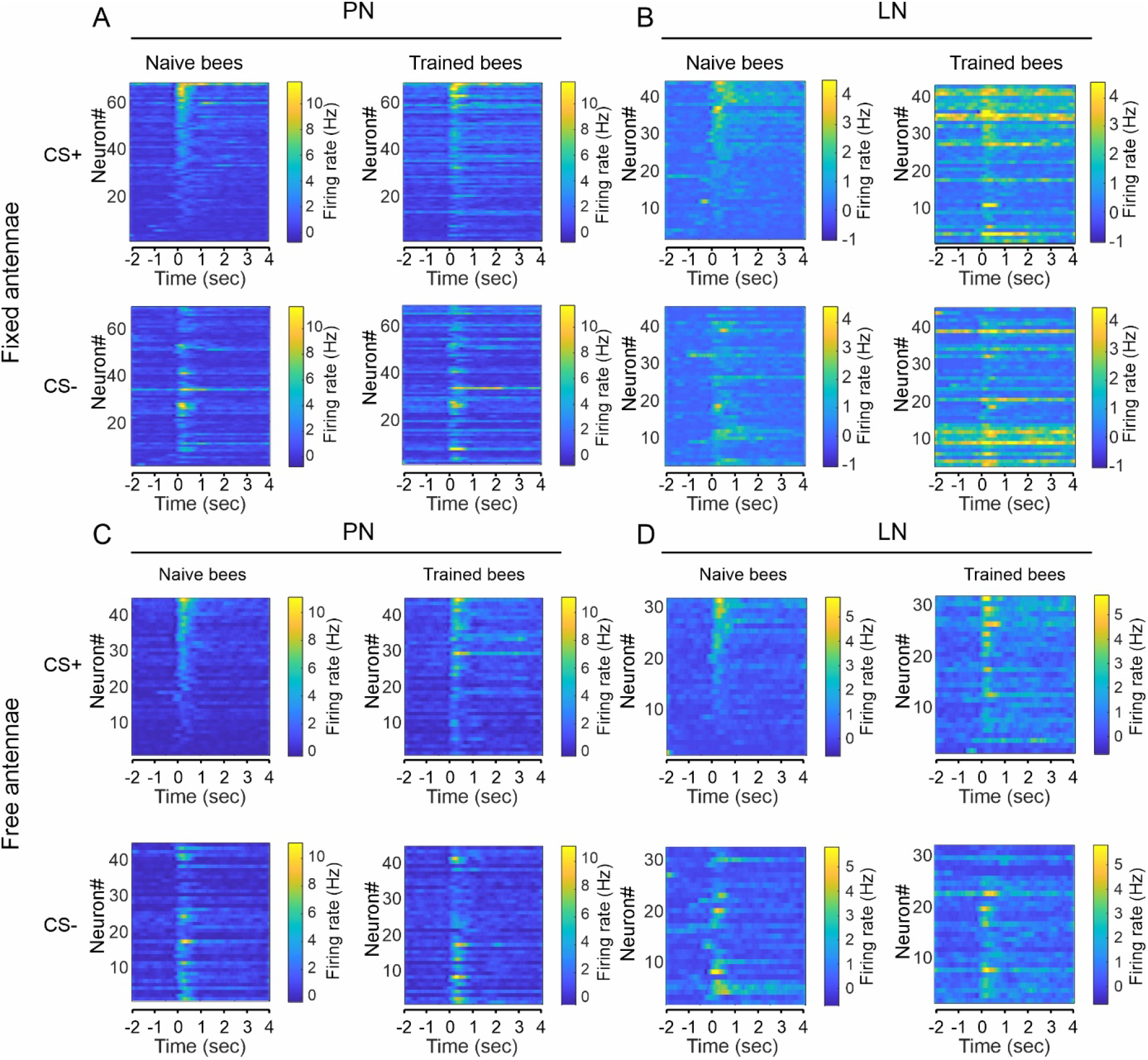
Peristimulus time histograms (psth) to odour stimulations (10^-1^ v/v dilution) under different conditions. The histograms were processed with a third-order Savitzky-Golay filter and sorted in descending order per response magnitude of naïve bees to CS+ odour. The four quadrants (A,B,C,D) show psth data from PNs and LNs of fixed- and free-antennae bees to CS+ and CS-odours. Odours were delivered between 0 and 1 sec. Individual PNs and LNs from all bees are vertically stacked in each heatmap panel. Note the scale of color bars for PN and LN is different, indicating the PNs produced stronger responses than the LNs. Overall, both PNs and LNs from the trained bees displayed elevated firing rate than the naïve bees.

The peristimulus time histograms (psth) of PNs and LNs displayed distinct response patterns to CS+ and CS-odours (Fig.2). In the bees with fixed antennae (Fig.2 A, B), before the differential training (i.e. naïve bees in A), some PNs responded more strongly to the CS+ odour (Fig.2A, neurons above #35), whereas the CS-odour elicited responses in a slightly overlapping but generally different set of neurons. After the differential training (i.e. trained bees in A) some PNs that did not respond to CS+ before training now responded to the CS+ odour (neuron# 1-30 in A). This change was also true with CS-stimulations. On the other hand, the LNs in the fixed group (Fig. 2B) generally showed a long-lasting response pattern, meaning the odour-evoked responses outlast the duration of odour presentation. Additionally, the training-induced elevation of firing rate was apparent in most LNs. In the free-antennae group (Fig. 2C, D), similar increases in firing rates with training occurred in both PNs and LNs.

To examine the potential effects of antenna treatments, we re-arranged the psth data in such a way that the averaged odour-evoked responses in the fixed- and free-antennae group could be directly compared (Fig.3). In the population responses of PNs (Fig.3A-D), the free-antennae treatment showed a tendency of increased firing at the peak of the response to the CS+ (Fig. 3 A, B) and CS- (Fig. 3 B, D) odours after training. However, the LNs from the free-antennae group had more pronounced significant responses to both CS+ (Fig. 3 E, F) and CS- (Fig. 3 G, H) odours relative to the fixed-antennae group. Even before conditioning, the LNs of naïve bees exhibited a tendency of stronger responses to both odours when antennae were free, suggesting a possible role of antenna movement in generating stronger responses in these neurons. Differences in LN responses in naïve bees became even more pronounced after training.

**Fig. 3.**
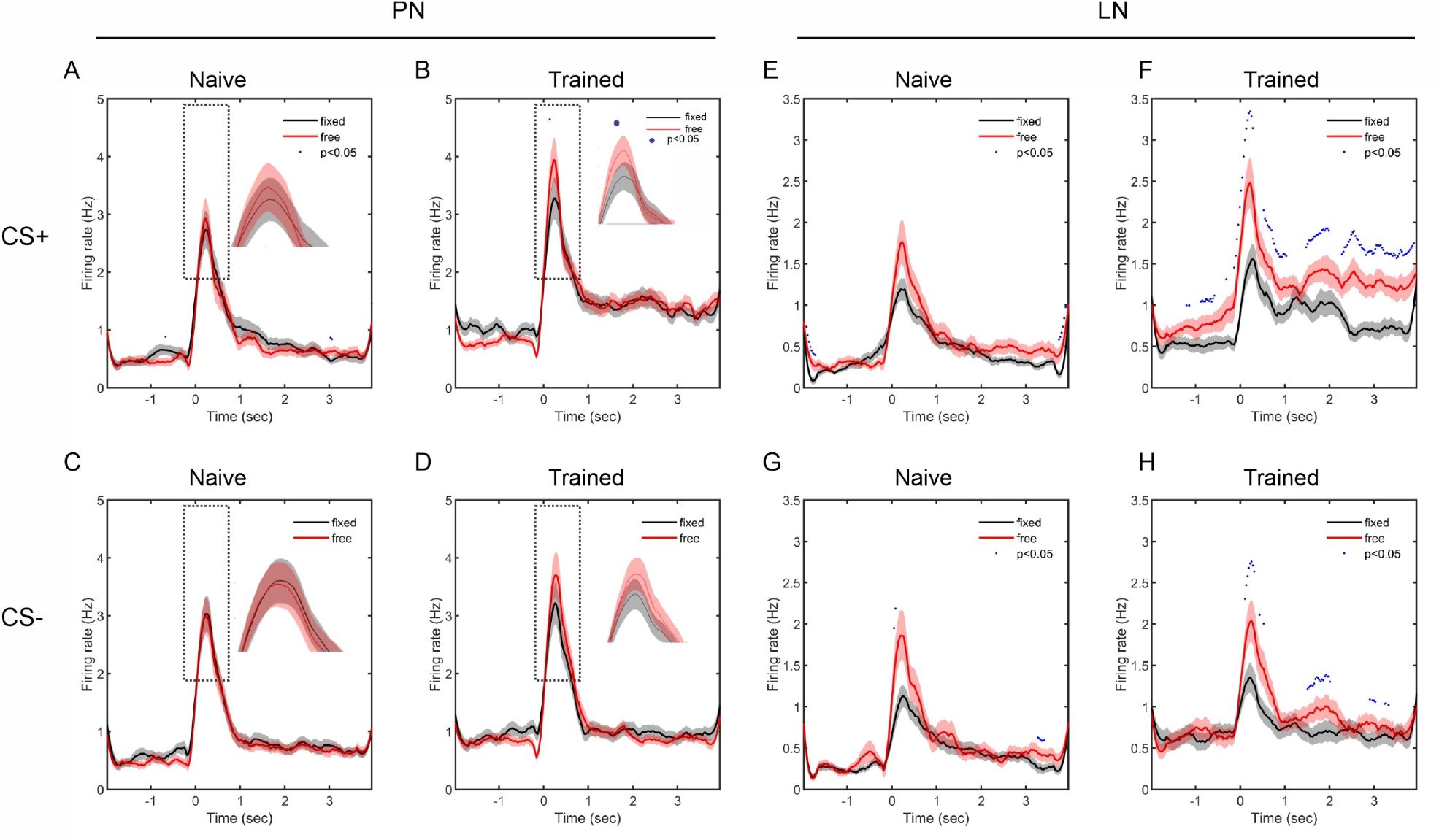
Variations of PN and LN response patterns to CS+ and CS-odour under different antenna treatments. In all panels, the shaded black curves represent the averaged peristimulus time histograms (psth) across all neurons (i.e. mean ± s.e.) in the fixed-antennae group, and the shaded red curves (mean ± s.e.) depict response patterns from the free-antennae group. Odour stimulations (10^-1^ vol/vol dilution) were delivered between 0 and 1 sec. The blue dots above the psth curves indicate statistical significance at those time points (Mann-Whitney U test, p<0.05). In PNs (A, B, C, D), the psth curves from the fixed- and free-antennae group are largely overlapping but the peak responses from the free-antennae group are higher than that from the fixed-antennae group after training (insets in B and D). In LNs (E, F, G, H), however, the free-antennae treatment resulted in significantly higher firing rate, especially after the training procedure.

Next, we evaluated the effect of antennal movement on plasticity across a broader range of concentrations given a 10^-1^ training concentration (Fig. 4). We quantified the response magnitude of PNs and LNs by calculating their averaged instantaneous firing rate within a defined response window (see Methods; Supplemental fig.1) for each concentration of CS+ and CS-odours under fixed- and free-antennae conditions. For the CS+ (Figs. 4A-D), both PNs and LNs tended toward increased firing rate with the increase of concentration, and, as in previous reports (Faber et al., 1999; Fernandez et al., 2009), the training procedure generated higher responses in both fixed- and free-antennae groups. However, the effects of concentration and training phase either were not significant or only barely reached significance (Table 1 main terms). Part of the reason for the weak or missing main effects was a set of significant interactions between test concentration and the other main effects (Supplemental Table 1 two-way interactions, i.e. antenna treatment : concentration (F=4.8 p <0.005); concentration : training phase (F=7.5 p<0.001); concentration : neuron (F=4.6 p<0.005)). The PNs (Fig.4. A, C) from the trained bees showed higher responses in all concentrations to the CS+ odour, but the increment with concentration is less pronounced in the fixed-antennae bees than in the free-antennae bees. The same interaction occurred in LNs, in which the differences across concentration were slightly greater than for PNs (Fig.4B versus 4D). The same training-induced effects were evident for the CS-odours (Fig. 4 E-H; Supplemental Table 2).

**Fig. 4.**
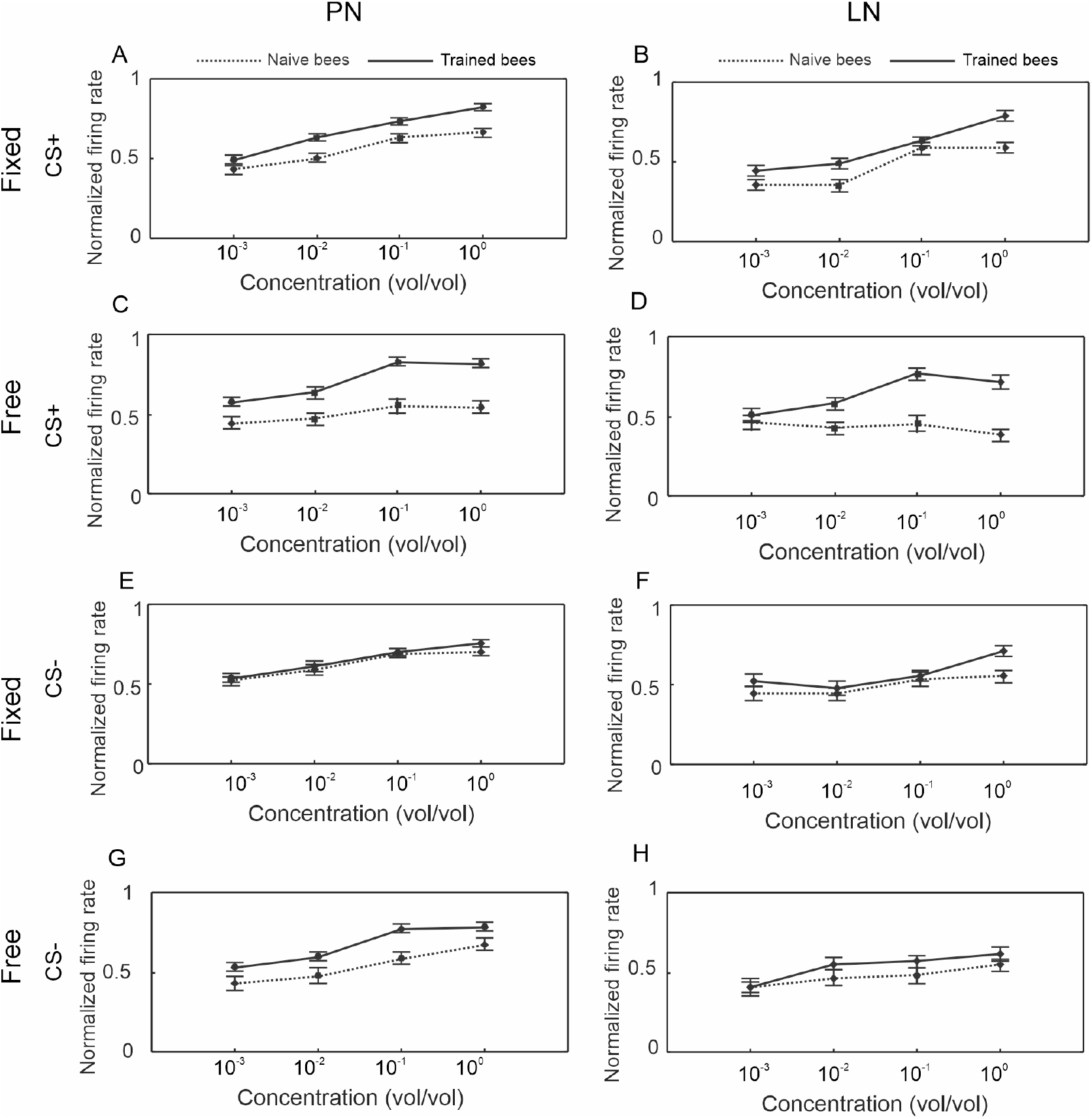
Dose-response curves of PNs (n=68) and LNs (n=44) to CS+ and CS-odours under free- or fixed-antennae conditions. The response values were normalized to the maximal response of each neuron across all four concentrations, both training phases, both odours and both antenna treatments, then the mean ± s.e. were used to construct the dose-response curves. The PN and LN data were processed separately. In each panel, the dotted line indicates the responses from naïve bees and the solid line indicates responses from trained bees. In PNs (A, C, E, G), the responses became stronger with increasing concentrations regardless of CS+ or CS-odour. The trained bees displayed elevated responses at all concentrations comparing with the naïve bees except for E (fixed, CS-). Comparison between A and C or B and D shows the difference between fixed- and free-antennae treatment.

To further illustrate how free antenna movement might have impacted training-induced changes in the antennal lobes, we plotted the response differences to CS+ and CS-odours between the trained bees and naïve bees from both the fixed- and the free-antennae groups (Fig.5). Overall, PNs from the free-antennae bees showed consistently larger changes to both CS+ and CS-odours across all four concentrations than PNs from the fixed-antennae bees with the largest increment at the 10^-1^ training concentration (Fig. 5 A, C), with the interaction of test concentration and antennal treatment being significant for both odours (Supplemental Table 3-4: CS+, F=4.6, p <0.005; CS-, F=9.7 p<0.001). The coding in the LNs was more variable, but with this analysis the effects of the free-antennae movement were still apparent in the CS+ stimulation at higher concentrations (Fig.5 B, D; Supplemental Table 5-6).

**Fig. 5.**
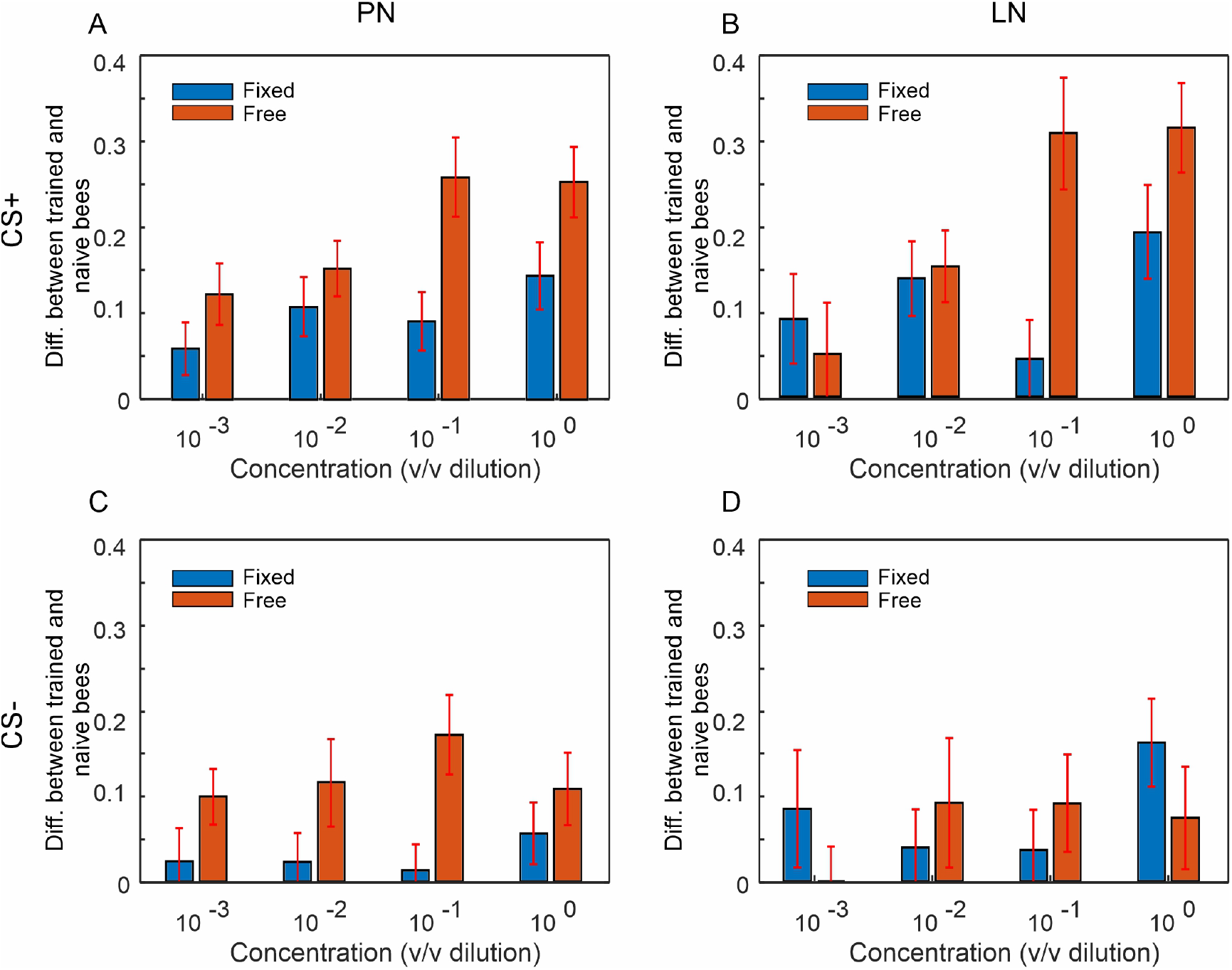
Effects of antenna treatments on associative learning. The increment of response magnitude as result of the differential training procedure, i.e. Response_trained_ - Response_naive_, is illustrated across all four concentrations (10^-3^, 10^-2^, 10^-1^, 10^0^ vol/vol dilutions) to CS+ (A, B) and CS- (C, D) odours. Normalized responses were quantified by the averaged instantaneous firing rate within the 1 sec response window (see Methods), and the differences between the response before training (i.e. naïve bees) and after training (i.e. trained bees) were calculated to show the training effects. In all panels, the training effects on the fixed-antennae bees and free-antennae bees are represented by blue bars (mean ± s.e.) and red bars (mean ± s.e.), respectively. In general, the free-antennae treatment resulted in stronger training effect (i.e. red bars are higher than blue bars), especially in PNs regardless of CS+ or CS-odours (A, C). In LNs (B, D), free-antennae treatment had less consistent effects.

## DISCUSSION

A growing body of work supports the interpretation that insect antennal movements can actively affect olfactory sensing (Cholé et al., 2022; Claverie et al., 2023; Hoffmann and Couzin-Fuchs, 2023; Huston et al., 2015; Lei et al., 2022). Antennal movements enhance peripheral and central processing of odours, presumably making them more salient. Yet to date very little evidence has shown how that peripheral or more central processing could be enhanced by antennal movement. Here we show when honey bee antennae are left free to move during an associative olfactory learning protocol, the result is a stronger engagement of the well-established plasticity in the antennal lobes, consistent with several previous studies that have shown that plasticity is important for separation of odour representations (Faber et al., 1999; Locatelli et al., 2016; Rath et al., 2011). We also show that the dose-response curve measured from freely moving antennae is similar to that when the antennae are fixed in place (Fig. 1 E), however, the responses are more variable particularly at lower concentrations (Fig. 1 F). This implies that the variation could be important for engaging the plasticity, and further implies that the enhanced plasticity is an important result of active sensing.

### Active antennal movements correlate with olfactory learning

The way that unrestrained antennae are moved depends on different contexts (Cholé et al., 2015; Cholé et al., 2022; Gascue et al., 2022; Jernigan et al., 2026), although for experiments reported here we simply allowed antennae to freely move without precise monitoring of movements. Therefore, a more thorough understanding of how movements affect plasticity will require more detailed correlations of the specific types of movements with neural activity. For example, when restrained for PER trials, honey bees move their antennae constantly at low frequency, and the position of the antennae alternates between two main positions relative to the airstream (Jernigan et al., 2026). Odour stimulation changes the frequency with which antennae adopt these positions. The mean position in the absence of odour is across and slightly upwind of the direction of airflow. Presentation of an odour, about which bees are naïve as to meaning, causes the mean position to move slightly downwind and directly across the airflow. After association of an odour with food reinforcement in the form of a sucrose-water droplet, bees reorient the frequency of antennal movements closer to upstream (toward the odour source). In that context movements are faster and contain higher frequency components involving low amplitude back-and-forth movements in the odour stream. The statistical pattern of these movements depends on the specific turbulent structure of the odour plume (Jernigan et al., 2026).

Hypothetically, variance of sensory inputs can be driven by the interactions of antennal movements with the complex, turbulent structure of an odour plume (Crimaldi and Koseff, 2001). Odour molecules diffuse from an odour source into an air flow, and turbulence then quickly breaks up the odour mass into filaments. The formation and dispersal of filaments are functions of air speed as well as friction generated by movement of air over the ground and around objects (Connor et al., 2018). These factors can all change the shape and encounter rates of odour filaments. Therefore, small changes in antennal position, velocity, or sweep geometry could alter whether and how often odour filaments intersect the antennal surface and how long they remain in contact with the antennae. Presumably these movements enhance interaction of antennae, and hence the olfactory sensory structures on them, with odour molecules carried in the flow.

Movements across the flow versus upwind toward the source could represent odour exploration versus exploitation (Jernigan et al., 2026), analogous to the caste and surge movements of insect flying in an odour plume (Stupski and van Breugel, 2024; Talley et al., 2023). Increasing the frequency that the antennae are in a cross-wind ‘exploratory’ position, which increases the cross-section of space that is sampled, might enhance the probability of filament encounter, particularly when filaments are sparse and infrequent. The orientation more upwind during ‘exploitation’, combined with high-frequency components, could increase exposure of sensory structures all along the antennae, and it could enhance penetration of molecules to those structures.

### Active antennal movements reshape the boundary layer on antenna

In insects, penetration of molecules at the antenna is constrained by near-surface transport processes (Goulet et al., 2025). Odorants must diffuse across a boundary layer before reaching olfactory sensilla, and boundary-layer dynamics can reshape the temporal and amplitudinal structures of the stimulus arriving at receptors (Schneider et al., 1998). Movement of air across the antennae, either when antennae are held relatively fixed in flight or are being moved by a stationary or walking bee, can shape how the boundary layer affects movement of molecules to sensory structures (Goulet et al., 2025). In honey bees, olfactory sensory neuron dendrites are housed in pore plates (Esslen and Kaissling, 1976; Frasnelli et al., 2010), which are oval depressions in the cuticle ringed with pores through which molecules gain access to hemolymph that bathes the dendrites. Pore plates are concentrated on the leading edge of the antennae, which faces upwind. Movement of air over the antennae changes the thickness of the boundary layer at the leading edge, which enhances odour capture at the pore plates (see figs 2 and 3 in (Goulet et al., 2025)).

We propose that antennal movement, particularly the way that it affects the boundary layer, is the cause for higher variance particularly at low concentrations when odour-carrying filaments are sparse. That is, antennal movement may induce variation at lower concentrations such that at times the input will be higher than the expected mean relative to the no movement condition. The variation could be important for detection of odours under natural conditions. In a recent review, Wachowiak et al (Wachowiak et al., 2025) noted that under natural conditions odour concentrations are typically only a few parts per billion, well under the concentrations typically used for laboratory-based investigations of olfaction. The lowest concentrations we use here would therefore be at the upper range for natural odours. This implies that active sensing antennal movements may be adapted for detection and encoding of odours under conditions bees would encounter while walking and flying in their natural environment.

In summary, our results show that active antennal movement modulates neural plasticity during olfactory learning. Bees learned to discriminate a rewarded odour (CS+) from a punished odour (CS−) equally well whether their antennae were freely movable or fixed in place (Supplemental figure 2). At the neuronal level, however, associative training generally increased firing rates in both projection neurons and local neurons, with the strongest enhancement occurring in responses to the CS+ (Fig. 4, 5). Importantly, bees with freely moving antennae exhibited a more pronounced differentiation between CS+ and CS− representations based on neuronal firing rate, suggesting that antennal movement enhances learning-related plasticity in the antennal lobe. One possible mechanism is the greater trial-to-trial variability in sensory responses observed when the antennae are free to move (Fig. 1), perhaps by reshaping air flow and boundary layer. Although this movement-enhanced plasticity does not appear necessary for initial acquisition of odour discrimination, it may influence the robustness, specificity, or accessibility of the resulting memory during later retrieval.

## ACKNOWLEDGEMENTS

This project is supported by the NSF/CIHR/FRQ/UKRI-MRC Next Generation Networks for Neuroscience Program (Award #2014217) to B.H.S and J.D.V. (www.odour2action.org). Also supported by National Institute of Health award (NIDCD R01 DC020892-01) to H.L..

## REFERENCES

Birgiolas, J., C.M. Jernigan, R.C. Gerkin, B.H. Smith, and S.M. Crook. 2017. SwarmSight: real-time tracking of insect antenna movements and proboscis extension reflex using a common preparation and conventional hardware. JoVE

Biswas, D., L.A. Arend, S.A. Stamper, B.P. Vágvölgyi, E.S. Fortune, and N.J. Cowan. 2018. Closed-Loop Control of Active Sensing Movements Regulates Sensory Slip. Curr. Biol. 28:4029-4036.e4024.

Cardé, R.T. 2021. Navigation Along Windborne Plumes of Pheromone and Resource-Linked Odours. Annual Review of Entomology 66:317–336.

Cholé, H., P. Junca, and J.-C. Sandoz. 2015. Appetitive but not aversive olfactory conditioning modifies antennal movements in honeybees. Learning & Memory 22:604–616.

Cholé, H., A. Merlin, N. Henderson, E. Paupy, P. Mahé, G. Arnold, and J.-C. Sandoz. 2022. Antenna movements as a function of odourants’ biological value in honeybees (Apis mellifera L.). Scientific reports 12:11674.

Claverie, N., P. Buvat, and J. Casas. 2023. Active Sensing in Bees Through Antennal Movements Is Independent of Odour Molecule. Integrative and Comparative Biology 63:315–331.

Connor, E.G., M.K. McHugh, and J.P. Crimaldi. 2018. Quantification of airborne odour plumes using planar laser-induced fluorescence. Experiments in Fluids 59:137.

Crimaldi, J.P., and J.R. Koseff. 2001. High-resolution measurements of the spatial and temporal scalar structure of a turbulent plume. Experiments in Fluids 31:90–102.

Esslen, J., and K.-E. Kaissling. 1976. Zahl und Verteilung antennaler Sensillen bei der Honigbiene (Apis mellifera L.). Zoomorphologie 83:227–251.

Faber, T., J. Joerges, and R. Menzel. 1999. Associative learning modifies neural representations of odours in the insect brain. Nature Neuroscience 2:74–78.

Fernandez, P.C., F.F. Locatelli, N. Person-Rennell, G. Deleo, and B.H. Smith. 2009. Associative Conditioning Tunes Transient Dynamics of Early Olfactory Processing. The Journal of Neuroscience 29:10191–10202.

Frasnelli, E., G. Anfora, F. Trona, F. Tessarolo, and G. Vallortigara. 2010. Morpho-functional asymmetry of the olfactory receptors of the honeybee (Apis mellifera). Behavioural brain research 209:221–225.

Gascue, F., E. Marachlian, M. Azcueta, F.F. Locatelli, and M. Klappenbach. 2022. Antennal movements can be used as behavioural readout of odour valence in honey bees. IBRO Neuroscience Reports 12:323–332.

Goulet, D., B. Smith, A. True, and J. Crimaldi. 2025. Pore plate sensilla scale and distribution modulate odour capture around honey bee antennae. Scientific reports 15:41574.

Hadar, R., and R. Menzel. 2010. Memory Formation in Reversal Learning of the Honeybee. Front. Behav. Neurosci. Volume 4 - 2010:

Hoffmann, A., and E. Couzin-Fuchs. 2023. Active smelling in the American cockroach. Journal of Experimental Biology 226:

Huston, S.J., M. Stopfer, S. Cassenaer, Zane N. Aldworth, and G. Laurent. 2015. Neural Encoding of Odours during Active Sampling and in Turbulent Plumes. Neuron 88:403–418.

Jernigan, C.M., E. Connor, H. Lei, J.D. Victor, J. Crimaldi, and B.H. Smith. 2026. Active sensing: different plume structures affect movements of antennae in honey bees (Apis mellifera). Journal of Experimental Biology 229:

Jernigan, C.M., R. Halby, R.C. Gerkin, I. Sinakevitch, F. Locatelli, and B.H. Smith. 2020. Experience-dependent tuning of early olfactory processing in the adult honey bee, Apis mellifera. The Journal of Experimental Biology 223:jeb206748.

Khan, A.G., M. Sarangi, and U.S. Bhalla. 2012. Rats track odour trails accurately using a multi-layered strategy with near-optimal sampling. Nature Communications 3:703.

Latshaw, J.S., R.E. Mazade, M. Petersen, J.A. Mustard, I. Sinakevitch, L. Wissler, X. Guo, C. Cook, H. Lei, J. Gadau, and B. Smith. 2023. Tyramine and its Amtyr1 receptor modulate attention in honey bees (Apis mellifera). eLife 12:e83348.

Lei, H., S. Haney, C.M. Jernigan, X. Guo, C.N. Cook, M. Bazhenov, and B.H. Smith. 2022. Novelty detection in early olfactory processing of the honey bee, Apis mellifera. PLOS ONE 17:e0265009.

Locatelli, F.F., P.C. Fernandez, and B.H. Smith. 2016. Learning about natural variation of odour mixtures enhances categorization in early olfactory processing. The Journal of Experimental Biology 219:2752.

Meyer, A., C.G. Galizia, and M.P. Nawrot. 2013. Local interneurons and projection neurons in the antennal lobe from a spiking point of view. Journal of Neurophysiology 110:2465–2474.

Murlis, J., J. Elikton, and R. Carde. 1992. Odour plumes and how insects use them. Annual Rev of Entomology 37:505–532.

Rath, L., C. Giovanni Galizia, and P. Szyszka. 2011. Multiple memory traces after associative learning in the honey bee antennal lobe. European Journal of Neuroscience no-no.

Sane, S.P., and N.P. Jacobson. 2006. Induced airflow in flying insects II. Measurement of induced flow. Journal of Experimental Biology 209:43–56.

Schneider, R.W.S., J. Lanzen, and P.A. Moore. 1998. Boundary-layer effect on chemical signal movement near the antennae of the sphinx moth, Manduca sexta: temporal filters for olfaction. Journal of Comparative Physiology A-Sensory Neural and Behavioural Physiology 182:287–298.

Smith, B.H., and C.M. Burden. 2014. A Proboscis Extension Response Protocol for Investigating Behavioural Plasticity in Insects: Application to Basic, Biomedical, and Agricultural Research. 1940-087X, e51057 pp.

Spencer, T.L., A. Clark, J. Fonollosa, E. Virot, and D.L. Hu. 2021. Sniffing speeds up chemical detection by controlling air-flows near sensors. Nature Communications 12:1232.

Stupski, S.D., and F. van Breugel. 2024. Wind gates olfaction-driven search states in free flight. Current Biology 34:4397-4411.e4396.

Suver, M.P., A.M. Medina, and K.I. Nagel. 2023. Active antennal movements in Drosophila can tune wind encoding. Current Biology 33:780-789.e784.

Talley, J.L., E.B. White, and M.A. Willis. 2023. A comparison of odour plume-tracking behaviour of walking and flying insects in different turbulent environments. Journal of Experimental Biology 226:

Thesen, A., J.B. Steen, and K.B. Døving. 1993. Behaviour of Dogs During Olfactory Tracking. Journal of Experimental Biology 180:247–251.

Wachowiak, M. 2011. All in a Sniff: Olfaction as a Model for Active Sensing. Neuron 71:962–973.

Wachowiak, M., A. Dewan, T. Bozza, T.F. O’Connell, and E.J. Hong. 2025. Recalibrating Olfactory Neuroscience to the Range of Naturally Occurring Odour Concentrations. The Journal of Neuroscience 45:e1872242024.

Zweifel, N.O., and M.J.Z. Hartmann. 2020. Defining “active sensing” through an analysis of sensing energetics: homeoactive and alloactive sensing. Journal of Neurophysiology 124:40–48.

