## supplemental figures for "Active sensing movements by antennae enhance neural processing and plasticity for odour processing in the CNS in the honey bee *Apis mellifera*"

SP fig.1

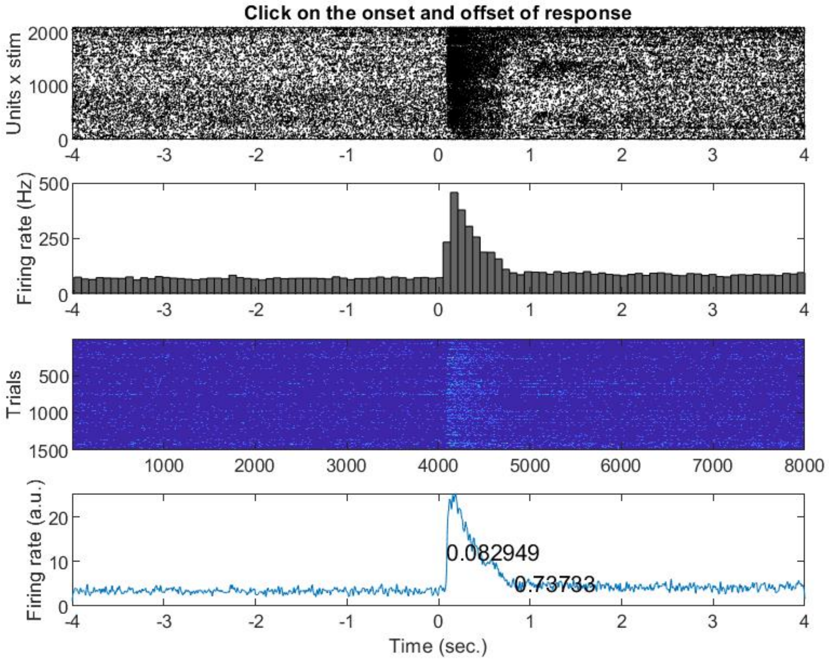

SP Fig.1 Data visualization for determining response window. This figure shows an example from the experiment labelled as NK20210521b. The response onset and offset are determined in Matlab, and subsequent calculations of response magnitude use the response window of 0.7373 sec -0.0829 sec = 0.6544 sec in this experiment.

SP fig.2

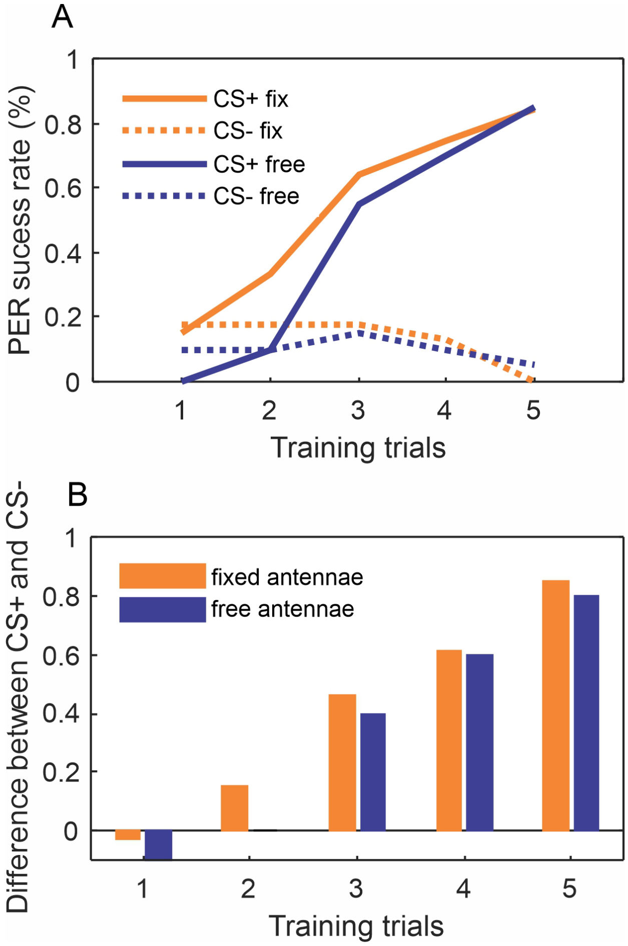

SP fig. 2 Behavioural response to differential training. Bees were subjected to 5 trials of differential conditioning with CS+ odour being rewarded with sugar solution and CS- odour being punished with salt solution. A positive behaviour featured proboscis extension reflex (PER) to conditioned odour. In both fixed-antennae group (solid orange line in A, n=39) and free-antennae group (solid blue line in A, n=20), CS+ elicited significantly more positive behaviours, compared with the CS- odour (dashed orange line versus dashed blue line in A). The difference between fixed- and free-antennae groups (two solid lines or two dashed lines in A), however, was not significantly different. This observation was further verified by calculating the difference between CS+ and CS- data, i.e. subtracting dashed line from solid line of same colour. The contrast between CS+ and CS- elicited PER rate becomes more pronounced over training trials (B). However, fixing antennae (blue bars in B) or keeping antennae free (orange bars in B) did not make significant difference in distinguishing CS+ and CS-.

SP.fig3

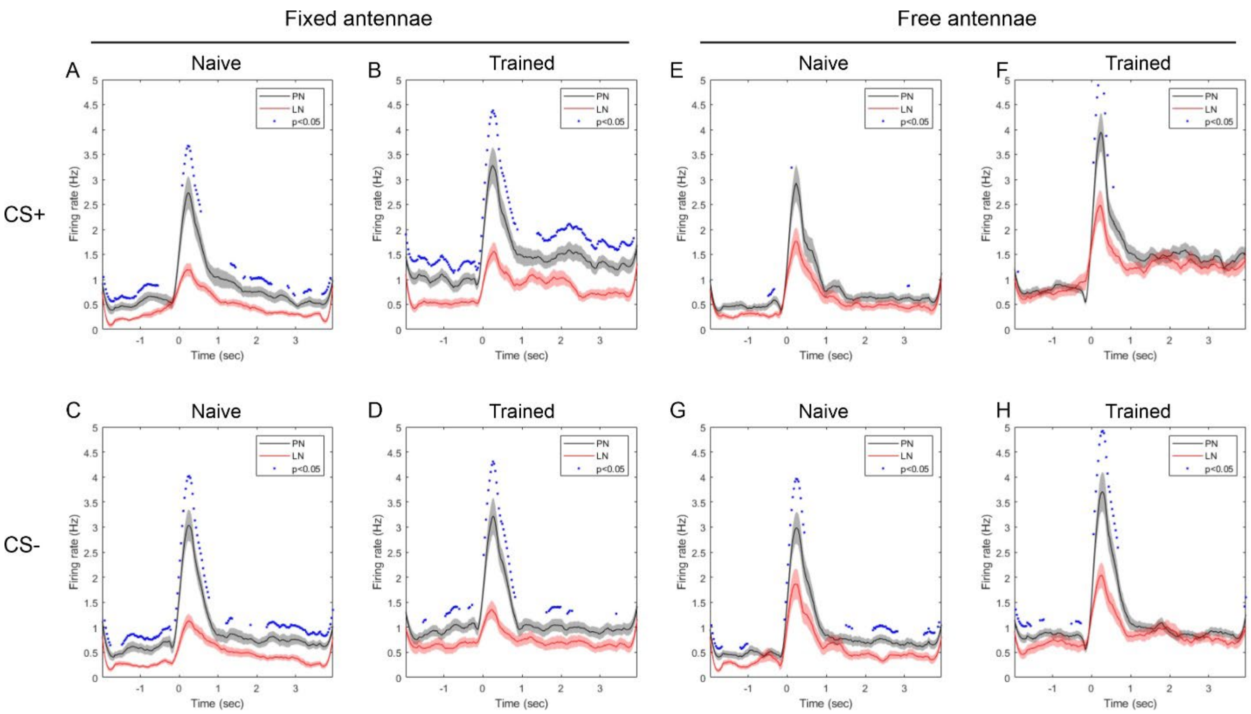

SP Fig.3 Responses of PNs and LNs to CS+ and CS- odours under fixed- and free-antennae conditions. The responses are shown as peristimulus time histograms (psth), which were constructed from spike data (binned at 10 ms) in a time window spanning 2 sec prior to and 4 sec after the onset of odour delivery executed with 5 pulses (pulse duration = 1 sec; interpulse interval = 30 sec). This time window is wide enough to cover all responses from both PNs and LNs. The histograms were then averaged from 68 PNs and 44 LNs, then filtered with a 3$^{rd}$-order Savitzky-Golay finite impulse response (FIR) filter with frame length of 21 points. In all panels, the odour onset was at 0 sec. The black curves (mean ± s.e.) represent the psth of PNs and the red curves (mean ± s.e.) were LNs. The blue dots mark significant differences between PN and LN at corresponding time points ($p<0.05$, Mann-Whitney U test). In both fixed- (A-D) and free-antennae group (E-H), the PNs had generally higher firing rate than LNs, confirming the validity of the method used for separating PNs and LNs in this study (Methods).

Supplemental Table 1. N-Way ANOVA on antennal treatment, odor concentration, training phase and neuron type for CS+ only.

| Source | Sum Sq. | d.f. | Mean Sq. | F | Prob>F |
|---|---|---|---|---|---|
| Treatment (fix vs free) | 0.003 | 1 | 0.00277 | 0.01 | 0.9136 |
| Concentration | 1.156 | 3 | 0.38525 | 1.64 | 0.1785 |
| Phase (pre vs post) | 0.654 | 1 | 0.65449 | 2.78 | 0.0954 |
| Neuron (PN vs LN) | 1.309 | 1 | 1.30921 | 5.57 | 0.0184 |
| Treatment (fix vs free):Concentration | 3.368 | 3 | 1.12271 | 4.78 | 0.0026 |
| Treatment (fix vs free):Phase (pre vs post) | 0.624 | 1 | 0.62381 | 2.65 | 0.1035 |
| Treatment (fix vs free):Neuron (PN vs LN) | 0.068 | 1 | 0.06827 | 0.29 | 0.59 |
| Concentration:Phase (pre vs post) | 5.255 | 3 | 1.75155 | 7.45 | 0.0001 |
| Concentration:Neuron (PN vs LN) | 3.223 | 3 | 1.07435 | 4.57 | 0.0034 |
| Phase (pre vs post):Neuron (PN vs LN) | 0.216 | 1 | 0.21568 | 0.92 | 0.3383 |
| Error | 307.686 | 1309 | 0.23505 | | |
| Total | 323.527 | 1327 | | | |

Supplemental Table 2. N-Way ANOVA on antennal treatment, odour concentration, training phase and neuron type for CS- only.

| Source | Sum Sq. | d.f. | Mean Sq. | F | Prob>F |
|---|---|---|---|---|---|
| Treatment (fix vs free) | 0.156 | 1 | 0.15561 | 0.67 | 0.4117 |
| Concentration | 1.756 | 3 | 0.58522 | 2.54 | 0.0552 |
| Phase (pre vs post) | 2.408 | 1 | 2.40827 | 10.44 | 0.0013 |
| Neuron (PN vs LN | 0.003 | 1 | 0.00303 | 0.01 | 0.9088 |
| Treatment (fix vs free):Concentration | 4.432 | 3 | 1.47718 | 6.4 | 0.0003 |
| Treatment (fix vs free):Phase (pre vs post) | 0.154 | 1 | 0.15368 | 0.67 | 0.4146 |
| Treatment (fix vs free):Neuron (PN vs LN | 0.061 | 1 | 0.06134 | 0.27 | 0.6062 |
| Concentration:Phase (pre vs post) | 6.398 | 3 | 2.13281 | 9.24 | 0 |
| Concentration:Neuron (PN vs LN | 5.735 | 3 | 1.91151 | 8.28 | 0 |
| Phase (pre vs post):Neuron (PN vs LN | 0.218 | 1 | 0.21795 | 0.94 | 0.3313 |
| Error | 379.632 | 1645 | 0.23078 | | |
| Total | 399.156 | 1663 | | | |

Supplemental table 3. Two-way ANOVA revealing the effects of training dependent on antennal treatments and odour concentrations in **PNs** on **CS+**. Data used in these analyses were the difference of responses between the trained and naïve bees. The main terms are antennal treatment and odour concentration.

| Source | Sum Sq. | d.f. | Mean Sq. | F | Prob>F |
|---|---|---|---|---|---|
| Treatment (fix vs free) | 72619.6 | 1 | 72619.6 | 7.59 | 0.0061 |
| Concentration | 88247.2 | 3 | 29415.7 | 3.08 | 0.0276 |
| Treatment (fix vs free):Concentration | 133269.7 | 3 | 44423.2 | 4.64 | 0.0033 |
| Error | 3673239.1 | 384 | 9565.7 | | |
| Total | 4015892.2 | 391 | | | |

Supplemental table 4. Two-way ANOVA revealing the effects of training dependent on antennal treatments and odour concentrations in **PNs** on **CS-**. Data used in these analyses were the difference of responses between the trained and naïve bees. The main terms are antennal treatment and odour concentration.

| Source | Sum Sq. | d.f. | Mean Sq. | F | Prob>F |
|---|---|---|---|---|---|
| Treatment (fix vs free) | 30.5 | 1 | 30.54 | 0.18 | 0.6748 |
| Concentration | 6419.3 | 3 | 2139.78 | 12.35 | 0 |
| Treatment (fix vs free):Concentration | 5018.4 | 3 | 1672.8 | 9.65 | 0 |
| Error | 85961.9 | 496 | 173.31 | | |
| Total | 99817.3 | 503 | | | |

Supplemental table 5. Two-way ANOVA revealing the effects of training dependent on antennal treatments and odour concentrations in **LNs** on **CS+**. Data used in these analyses were the difference of responses between the trained and naïve bees. The main terms are antennal treatment and odour concentration.

| Source | Sum Sq. | d.f. | Mean Sq. | F | Prob>F |
|---|---|---|---|---|---|
| Treatment (fix vs free) | 361 | 1 | 360.979 | 5.08 | 0.0251 |
| Concentration | 1778.6 | 3 | 592.856 | 8.34 | 0 |
| Treatment (fix vs free):Concentration | 469.2 | 3 | 156.394 | 2.2 | 0.0885 |
| Error | 18772.5 | 264 | 71.108 | | |
| Total | 21461.9 | 271 | | | |

Supplemental table 6. Two-way ANOVA revealing the effects of training dependent on antennal treatments and odour concentrations in **LNs** on **CS-**. Data used in these analyses were the difference of responses between the trained and naïve bees. The main terms are antennal treatment and odour concentration.

| Source | Sum Sq. | d.f. | Mean Sq. | F | Prob>F |
|---|---|---|---|---|---|
| Treatment (fix vs free) | 273.5 | 1 | 273.46 | 2.59 | 0.1083 |
| Concentration | 3135.9 | 3 | 1045.31 | 9.91 | 0 |
| Treatment (fix vs free):Concentration | 549 | 3 | 183 | 1.74 | 0.1596 |
| Error | 33739.7 | 320 | 105.44 | | |
| Total | 38468.1 | 327 | | | |
